# Impact of efavirenz, ritonavir-boosted lopinavir and nevirapine based antiretroviral regimens on the pharmacokinetics of lumefantrine and safety of artemether-lumefantrine in *falciparum*-negative HIV-infected Malawian adults stabilized on antiretroviral therapy

**DOI:** 10.1101/337410

**Authors:** Clifford George Banda, Fraction Dzinjalamala, Mavuto Mukaka, Jane Mallewa, Victor Maiden, Dianne J Terlouw, David G. Lalloo, Saye H. Khoo, Victor Mwapasa

## Abstract

There is conflicting evidence of the impact of commonly used antiretroviral therapies (ARTs) on the pharmacokinetics of lumefantrine and safety profile of artemether-lumefantrine. We compared the area under the concentration-time curve (AUC_0-14 days_) of lumefantrine and safety profile of artemether-lumefantrine in malaria-negative human immunodeficiency virus (HIV) infected adults in two steps. In step 1, a half-dose adult course of artemether-lumefantrine was administered as a safety check in four groups (n = 6/group): (i) antiretroviral-naïve, (ii) on nevirapine-based ART, (iii) on efavirenz-based ART and (iv) on ritonavir-boosted lopinavir-based ART. In step 2, a standard-dose adult course of artemether-lumefantrine was administered to a different cohort in three groups (n = 10–15/group): (i) antiretroviral-naïve, (ii) on efavirenz-based ART and (iii) on ritonavir-boosted lopinavir-based ART. In step 1, lumefantrine’s AUC_0-14 days_ was 53% [95% CI: 0.27-0.82] lower in the efavirenz-based ART group than the ART-naïve group and was 2.4 [95% CI: 1.58-3.62] and 2.9 [95% CI: 1.75-4.72] times higher in the nevirapine and ritonavir-boosted lopinavir groups, respectively. In step 2, lumefantrine’s AUC_0-14 days_ was 1.9 [95% CI: 1.26-3.00] times higher in the ritonavir-boosted lopinavir group and not significantly different between the efavirenz-and ART-naïve groups (0.99 [95% CI: 0.63-1.57]). Frequent cases of haematological abnormalities (thrombocytopenia and neutropenia) were observed in the nevirapine group in step 1, leading to a recommendation from the data and safety monitoring board not to include a nevirapine group in step 2. Artemether-lumefantrine was well tolerated in the other groups. The therapeutic implications of these findings need to be evaluated among HIV-malaria co-infected adults.

## INTRODUCTION

In sub-Saharan Africa (SSA), human immunodeficiency virus (HIV) and *Plasmodium falciparum (Pf)* malaria infections are co-endemic. HIV infection increases susceptibility to malaria (1–3), severity of *Pf* malaria and reduces the efficacy of some antimalarial drugs (4, 5). To combat these infections, the WHO recommends initiation of antiretroviral therapy (ART) in HIV-positive (HIV +) individuals regardless of their CD4 cell counts (6) and prompt use of artemisinin-based combination therapies (ACTs) for malaria infected individuals (7). The most commonly used ART in SSA contain non-nucleoside reverse transcriptase inhibitors (NNRTIs) such as efavirenz (EFV) and nevirapine (NVP) or protease inhibitors (PIs) such as ritonavir-boosted lopinavir (LPV/r). Artemether-lumefantrine (AL) is the most widely implemented first line ACT in the SSA region (3). HIV-malaria co-infection is common in SSA hence a large number of HIV + people on ART require concurrent treatment with AL.

Pharmacokinetic (PK) interactions between NNRTI or PI-containing ART and ACTs are likely since these classes of drugs affect the activity of cytochrome-P(CYP) 450 liver enzymes, including CYP3A4 and CYP2B6 (8–11). The interactions may impact the longer acting partner drug of an ACT which is vital in preventing post-treatment malaria recrudescence, after the rapid elimination of the artemisinins (12). Previous PK studies have found lower lumefantrine levels in healthy volunteers co-treated with AL and EFV-based ART and higher lumefantrine levels in those co-treated with AL and LPV/r-based ART when compared to those treated with AL only (13–15). However, PK studies on AL and NVP-based ART, have produced conflicting results, with some finding higher, lower or similar lumefantrine levels in HIV + individuals on NVP-based ART than ART-naive individuals treated with AL only (16–20). Furthermore, few studies have reported the safety profiles of co-administering AL with commonly used antiretroviral drugs in HIV infected individuals stabilized on ART.

To further characterize the impact of nevirapine-, efavirenz-, or ritonavir-boosted lopinavir-based ART on the PK of lumefantrine and safety profile of AL, we conducted an intensive PK study to compare secondary PK parameters of lumefantrine and the incidence of treatment-emergent adverse events in malaria-negative HIV-infected adults taking AL plus NVP-, EFV-, or LPV/r-based ART or AL only.

## MATERIALS AND METHODS

### Study design

An open-label, sequential group, PK study was conducted from August 2010 to March 2013 at Queen Elizabeth Central Hospital, in Blantyre, Malawi. The study was implemented in the following two steps: In step 1 [PACTR2010030001871293], a half adult dose of the AL (2 tablets of AL [Coartem®, Novartis] each tablet containing 20mg/120mg, artemether/lumefantrine) was administered at times 0, 8, 24, 36, 48 and 60 hours to malaria-negative HIV + individuals in the following groups: 1) an antiretroviral naive (control) group, and those receiving 2) NVP-based ART, 3) EFV-based ART and 4) LPV/r-based ART. This step served mainly as a preliminary safety evaluation, checking for unexpected clinical toxicities or interactions.

In step 2 [PACTR2010030001971409], after review of safety data from step 1 by an independent data and safety monitoring board (DSMB), a full standard dose of AL (4 tablets of Coartem^®^, each tablet containing 20mg/120mg AL) was administered at times 0, 8, 24, 36, 48 and 60 hours to a separate cohort of malaria-negative HIV + individuals in the following groups: 1) an antiretroviral naive (control) group, and those receiving 2) EFV-based ART and 3) LPV/r-based ART. The DSMB recommended that step 2 should not proceed with a NVP-based ART group because of safety concerns.

To maximize the absorption of lumefantrine, AL was given with ∽40 millilitres of soya milk, containing an equivalent of 1.2g of fat. The first dose of AL in ART participants was timed to coincide with the next scheduled dose of the antiretroviral drugs.

### Study population

The study population for step 1 and step 2 were HIV infected male and non-pregnant female adults aged ≥18 years residing in Blantyre, Malawi or neighbouring districts of Thyolo and Chiradzulu. Individuals on ART were eligible to participate if they had been on NNRTI or PI-based ART for ≥ 6 months and had CD4 cell count ≥ 250 cells/mm^3^. At the beginning of the study, HIV infected antiretroviral naive individuals were eligible if they had CD4 cell count ≥ 250/mm^3^ but this cut-off point was changed to ≥350/mm^3^ when the WHO criteria for ART initiation changed in July 2011. Other inclusion criteria were body weight ≥40kgs, willingness to be admitted in the hospital for 3 days, to remain within the study sites and to be contacted at home or by phone during the course of the study.

We excluded subjects who had a body mass index of <18.5kg/m^2^, haemoglobin concentration of <10 g/dL (subsequently changed to <8.5 g/dL based on DSMB recommendation), reported use of any antimalarial drugs within the preceding 4 weeks, reported hypersensitivity to any of the ACTs, receipt of other drugs which are known inhibitors or inducers of P450 enzymes or P-glycoprotein (except cotrimoxazole prophylaxis which was standard of care for HIV infected individuals), a history of regular intake of alcohol (>twice/week), tobacco (>3 times/week) or any use of illicit drugs, history or evidence of pre-existing liver, kidney or heart disease, including conductive abnormalities on electrocardiographs (QTc interval>450ms in men and >470ms in females), clinical and/or laboratory evidence of *Pf* malaria, hepatitis B, pneumonia, tuberculosis, bacteremia, laboratory evidence of potentially life threatening white blood cell disorders such as absolute neutrophil count <0.500*10^9^/L, absolute lymphocyte count <0.35*10^9^/L or absolute platelet count <25*10^9^/L, Karnofsky score of <80% or concurrent participation in any other clinical trial.

### Sample size

The sample size in step 1 was 6 in each of the AL/ART and control (ART-naive) and this was based on standard practice in early PK studies of antimalarial drugs which aim to safeguard the safety of study subjects and minimize the number of subjects who may be potentially exposed to harmful drug levels. In step 2, the sample size was 15 per group which had at least 90% power to detect a two-fold increase in lumefantrine AUC in any of the AL/ART groups compared with the ART-naive group, assuming a mean (standard deviation) lumefantrine AUC of 0.561 (0.36) μg/mL/hr (15) in the ART-naive group, at the level of significance of 5%.

### Ethics and screening procedures

The design and timing of trial procedures was approved by the College of Medicine Research Ethics Committee (COMREC), in Blantyre, Malawi. The study conformed to the principles of the International Conference on Harmonization on Good Clinical Practice. Research nurses and clinicians sought written informed consent from individuals to perform screening procedures including physical, medical and anthropometric assessment, electrocardiographs (ECGs) and blood tests to detect blood-borne infections, haematological, renal or hepatic abnormalities. Results from screening procedures were available within seven days of screening. Based on these results, potential study participants were informed of their eligibility to participate in the study. Thereafter, research nurses or clinicians sought written informed consent from eligible subjects to participate in the study.

### Pre-dosing procedures

Consenting study participants were re-assessed by research nurses or clinicians to determine whether they still met all eligibility criteria, through repeat history taking and physical examination. Eligible participants were admitted in hospital and an indwelling cannula was inserted into a vein before their scheduled dose of ART and the first dose of the ACT. Approximately 1 hour before the scheduled time of ART and ACT dosing, blood samples were collected for haematological, renal and liver function tests and random glucose test.

### Blood sample collection and processing

During hospitalization, blood samples for PK assays were collected in heparin tubes before treatment and at 1, 2, 4, 6, 12, 24, 36, 48, 60 and 72 hours post treatment. After discharge, blood samples were taken at 4, 5, 6, 7 and 14 days. Immediately after collection, the blood samples were spun in a refrigerated centrifuge and the separated plasma samples were temporarily frozen in liquid nitrogen before being transferred to a-80°C freezer until PK analyses.

### Safety assessments

After the first dose of AL, blood samples were collected to detect haematological, renal and liver function abnormalities at 12, 48 and 72 hrs and on day 7, 14, 21 and 28. In addition, in step 2, 12-lead electrocardiographs were performed pre-dosing, 5 hours after the first dose and 5 hours after the last dose to determine QTc interval using the Fridericia QT correction formula (21). The study focussed on treatment emergent adverse events (AEs) defined as “any clinical or subclinical abnormalities which were absent before dosing with AL but emerged post dosing or those which were present before dosing with AL but worsened post-dosing”. Severity of AEs was graded using the DAIDS criteria (22) while seriousness was defined according to the standard definition.

### Pharmacokinetic assays

Plasma samples were analysed for lumefantrine levels at the Malawi-Liverpool Wellcome Trust Clinical Research Programme in Blantyre, Malawi, using a validated HPLC-UV assay adopted and transferred to Malawi from Liverpool School of Tropical Medicine. The PK laboratory in Blantyre participated in WWARN’s External Quality Assurance programme (23). Briefly, lumefantrine and the internal standard (IS, Halofantrine), were recovered from plasma using a single protein precipitation step with acetonitrile and acetic acid (99:1). The supernatant was then evaporated to dryness in a vacuum concentrator at 25 ^°^C. The dried extract was re-dissolved in the reconstitution solvent methanol: hydrochloric acid 0.01M (70:30) and 75 μL injected into the chromatograph (Agilent 1100). Quantitation of the drugs was achieved by reverse phase HPLC. The optimum detection wavelength for each drug was 335 nm. The lower limit of quantification (LLQ) of the HPLC-UV assay was 0.05 μg/mL for lumefantrine with % coefficient of variation of <10. Extracted plasma pharmacokinetic samples were run in batches comprising of all samples collected from each of any two study participants. Each batch run included a blank plasma extract, two sets of 8-concentration-level calibration standards, and quality controls (QC) at three concentration levels: low, medium and high (0.05, 10 and 15 μg/mL). For batch assay to pass, the measured concentrations of at least 67% of the QC samples had to be within + /-20% of their nominal value and at least one QC had to be acceptable at the LLQ. The mean inter-assay precision for low, medium and high QCs was 6.6%, 8.8% and 9.2% respectively. In addition, 75% of each calibration curve’s concentrations had to lie within + /-20% and + /-15% of the nominal concentration at the LLQ or all other concentrations, respectively.

### Data analyses

Plasma concentrations of lumefantrine were analysed using non-compartmental pharmacokinetic analysis (NCA), employing the trapezoidal rule with cubic splines. Observed lumefantrine concentrations below the lower limit of quantification (<LLOQ) were treated as missing data except for the pre-dose lumefantrine concentration which was imputed to 0 if below LLOQ. For each study participant, the following PK parameters were computed: AUC_0-14 days_, maximum concentration [C_max_], time to maximum concentration [t_max_] and terminal elimination half-life [t_1/2_]). We used STATA 15.0 for the NCA and to compare log-transformed PK parameters. Geometric mean ratios with 95% confidence intervals have been presented. To test for significant differences in PK parameters between each ACT/ART group and the ART-naïve group, parametric evaluation of the log-transformed PK parameters was done using analysis of variance (ANOVA) (α = 0.05). Fisher’s exact test was used to compare proportions of participants across the study groups with day 7 concentrations that were above a value known to predict treatment response by day 28, and of safety parameters across the different ACT/ART groups in comparison to the ART naïve group. Data summaries and graphics were all performed in STATA 15.0.

## RESULTS

### Characteristics of participants

In step 1, 26 participants were enrolled in the study; 24 participants were successfully followed up for 28 days. Two participants taking NVP-based ART were discontinued from the study due to protocol deviations and are not included in the analyses. In step 2, 40 of the 43 enrolled study participants completed 28 days of follow-up. Three participants did not have sufficient data points for PK characterization and are not included in the analyses. No participants were enrolled in the NVP arm for step 2 on the advice of the DSMB because of the observed haematological abnormalities in step 1. Supplementary Table 1 shows the baseline characteristics of participants who completed follow-up in steps 1 and 2. In step 1, the median duration of ART (in months) was significantly longer in the LPV/r group (63.1, range [33.3-85.0]) than in the EFV group (25.1, range [7.8-49.3]) and the NVP group (58.8, range [24.7-80.6]). There were no major differences between baseline characteristics in step 1 or step 2.

**Table 1:**
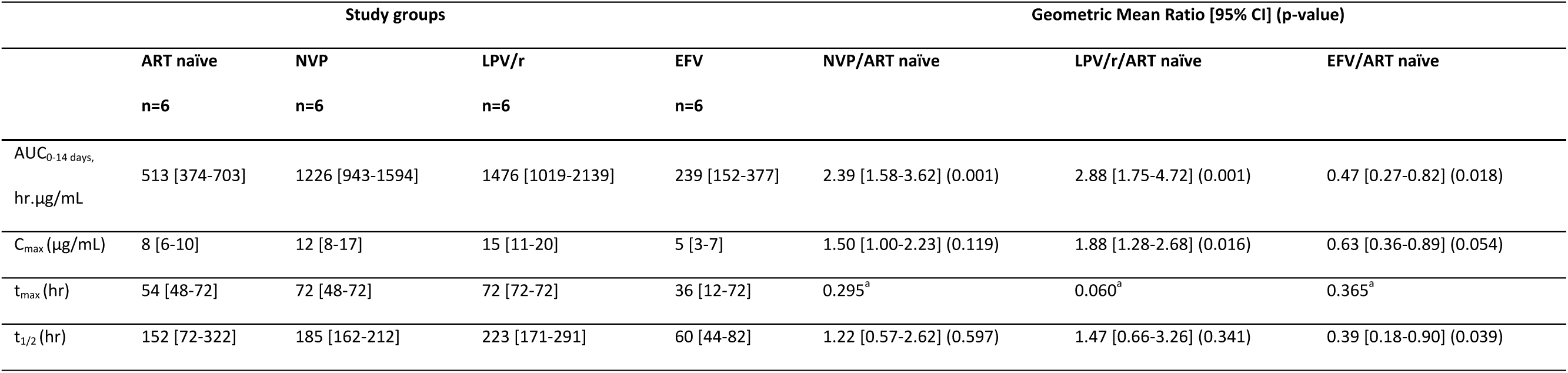
Lumefantrine pharmacokinetic parameters for participants in step 1

|  | Study groups |  |  |  | Geometric Mean Ratio [95% CI] (p-value) |  |  |
| --- | --- | --- | --- | --- | --- | --- | --- |
|  | ART naïve | NVP | LPV/r | EFV | NVP/ART naïve | LPV/r/ART naïve | EFV/ART naïve |
|  | n=6 | n=6 | n=6 | n=6 |  |  |  |
| AUC<sub>0-14 days</sub>,<br>hr.µg/mL | 513 [374-703] | 1226 [943-1594] | 1476 [1019-2139] | 239 [152-377] | 2.39 [1.58-3.62] (0.001) | 2.88 [1.75-4.72] (0.001) | 0.47 [0.27-0.82] (0.018) |
| C<sub>max</sub> (µg/mL) | 8 [6-10] | 12 [8-17] | 15 [11-20] | 5 [3-7] | 1.50 [1.00-2.23] (0.119) | 1.88 [1.28-2.68] (0.016) | 0.63 [0.36-0.89] (0.054) |
| t<sub>max</sub> (hr) | 54 [48-72] | 72 [48-72] | 72 [72-72] | 36 [12-72] | 0.295<sup>a</sup> | 0.060<sup>a</sup> | 0.365<sup>a</sup> |
| t<sub>1/2</sub> (hr) | 152 [72-322] | 185 [162-212] | 223 [171-291] | 60 [44-82] | 1.22 [0.57-2.62] (0.597) | 1.47 [0.66-3.26] (0.341) | 0.39 [0.18-0.90] (0.039) |

### Pharmacokinetics of lumefantrine and interactions with antiretroviral therapy in step 1

Table 1 summarizes the PK parameters in the study groups in step 1. Compared with the ART-naïve group, the geometric mean AUC_0-14 days_ of lumefantrine was 53% lower in the EFV-ART group, 2.4 times higher in the NVP-ART group and 2.9 times higher in the LPV/r-based ART group. Similarly, compared with the ART naïve group, lumefantrine’s C_max_ was 37% lower in the EFV-ART group (marginally significant), 1.9 times higher in the LPV/r-ART group and not significantly different in the NVP based ART arm. Additionally, compared with the ART naïve group, lumefantrine’s terminal half-life was 61% shorter in the EFV-group but not significantly different in the LPV/r-based and NVP-based ART groups. The median t_max_ was similar in the NVP-, EFV-based and ART-naïve groups but slightly longer in the LPV/r-based ART group than in the ART naïve group with marginal significance. As illustrated in the concentration-time profile in Figure 1a, participants in the LPV/r-and NVP-ART groups had higher concentrations of lumefantrine in the terminal elimination phase than those in the ART naïve sub-group, while those in the EFV-based ART group had lower lumefantrine concentrations.

**Figure 1a.**
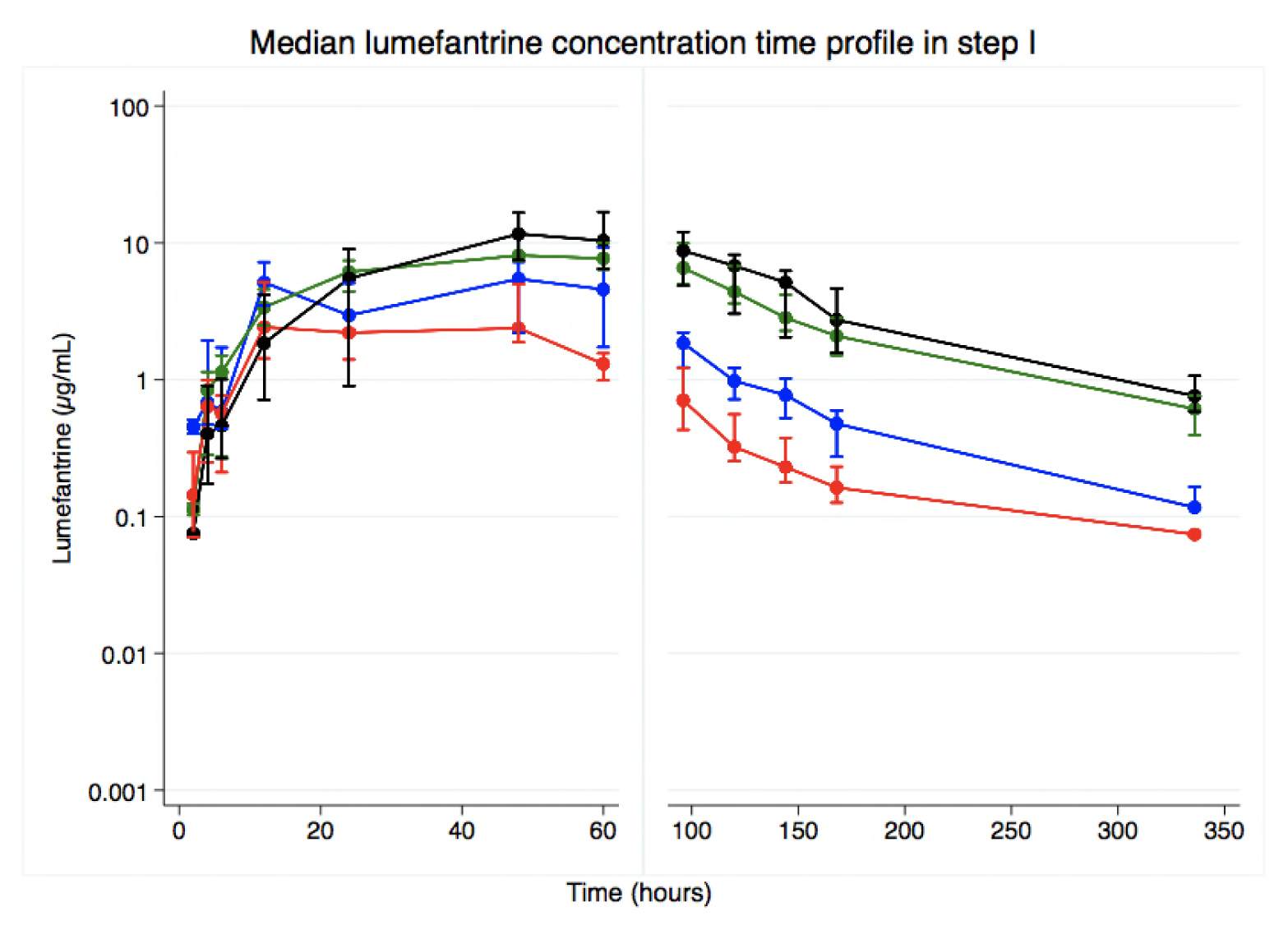
Plasma lumefantrine concentration-time profile in step 1 following administration of half (n = 24) adult treatment course of artemether-lumefantrine among antiretroviral therapy naïve (blue), those on efavirenz-(red), nevirapine-(green), and ritonavir-boosted lopinavir-(black) based antiretroviral therapy. Data are presented as median (IQR).

### Tolerability and safety of artemether-lumefantrine in step 1

AL was well tolerated in all the groups. However, DAIDS grade 3 or 4 treatment-emergent neutropenia were frequently detected across all the study groups: ART-naïve (3/6 [50.0%]), EFV-based ART (1/6 [16.7%]), LPV/r-based ART (2/6 [33.3%]) and NVP-based ART (3/6 [50.0%]). The inter-group differences were not statistically significant. Additionally, DAIDS grade 3 or 4 treatment-emergent thrombocytopenia was detected in the NVP-based ART (2/6 [33.3%]) but not in the ART-naïve or the LPV/r-and EFV-based ART groups. None of these observed adverse events were persistent beyond day 14 of follow up.

### Pharmacokinetics of lumefantrine and interactions with antiretroviral therapy in step 2

Table 2 summarizes the PK parameters in the study groups in step 2. The geometric mean lumefantrine AUC_0-14 days_ was similar in the EFV-based ART group and the ART-naïve group. Participants in the LPV/r-based ART group had an approximately 1.9 times higher geometric mean AUC_0-14 days_ than those in the ART naïve group. There were no significant differences in C_max_, t_1/2_ and median t_max_ in the EFV-and LPV/r-based ART groups compared to the ART-naïve group. As illustrated in the concentration-time profile in Figure 1b, lumefantrine concentrations were higher in the LPV/r-based ART than the ART-naïve group and were persistently lower in the terminal elimination phase (after 72 hours) in the EFV-based ART group than the ART-naïve group.

**Figure 1b.**
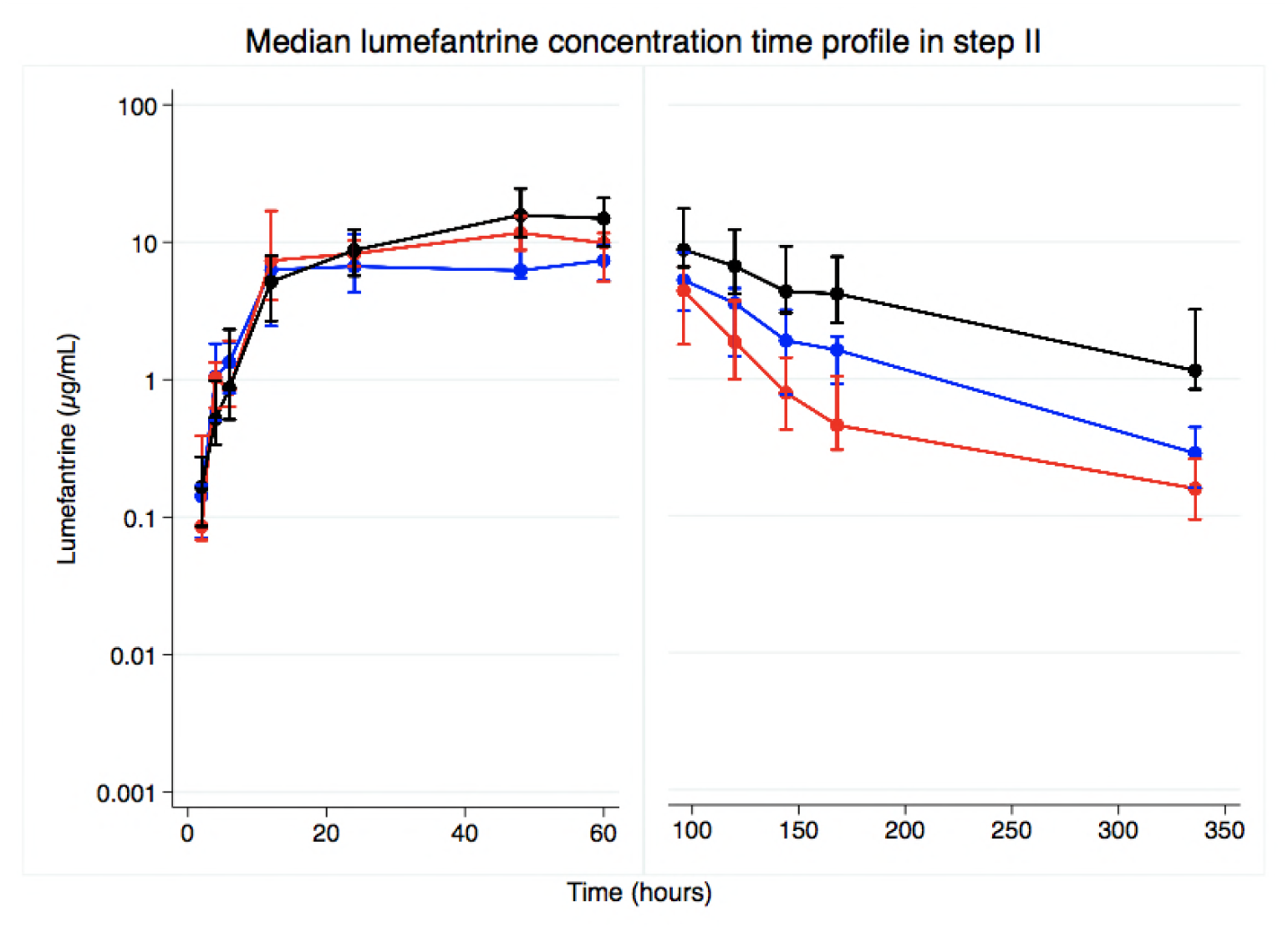
Plasma lumefantrine concentration-time profile in step 2 following administration of full-adult treatment course (n = 40) of artemether-lumefantrine among antiretroviral therapy naïve (blue), those on efavirenz-(red) and ritonavir-boosted lopinavir-(black) based antiretroviral therapy. Data are presented as median (IQR).

**Table 2:** Lumefantrine pharmacokinetic parameters for participants in step 2

|  | Study groups |  |  | Geometric Mean Ratio [95% CI] (p-value) |  |
| --- | --- | --- | --- | --- | --- |
|  | ART naïve<br>n=10 | LPV/r<br>n=15 | EFV<br>n=15 | LPV/r/ART naïve | EFV/ART naïve |
| AUC<sub>0-14 days</sub>,<br>hr.µg/mL | 1084 [760-1547] | 2107 [1654-2686] | 1081 [816-1432] | 1.94 [1.26-3.00] (0.004) | 0.99 [0.63-1.57] (0.991) |
| C<sub>max</sub> (µg/mL) | 15 [10-23] | 19 [16-23] | 18 [14-23] | 1.27 [0.81-1.93] (0.265) | 1.20 [0.75-1.84] (0.456) |
| t<sub>max</sub> (hr) | 66 [24-72] | 72 [60-72] | 48 [12-72] | 0.145<sup>a</sup> | 0.340<sup>a</sup> |
| t<sub>1/2</sub> (hr)* | 160 [103-248] | 190 [154-236] | 102 [61-170] | 1.19 [0.73-1.94] (0.438) | 0.64 [0.32-1.26] (0.217) |
| C<sub>d7</sub> (µg/mL) | 1 [0.9-2] | 4 [3-6] | 0.5 [0.3-0.8] | 4.00 [1.72-5.39] (<0.001) | 0.50 [0.21-0.74] (0.009) |

### Day 7 lumefantrine concentrations in step 2

Upon administration of a full standard AL dose, day 7 mean lumefantrine was 50% lower in the EFV-based ART group than in the ART-naïve group. Participants in the LPV/r-based ART group had 4 times higher day 7 lumefantrine concentration compared to those in the ART-naïve group as shown in Table 2. However, the proportion of participants with day 7 lumefantrine concentrations ≥0.2 μg/mL (200 ng/mL) were not significantly different in the ART-naïve group (100% [10/10]), LPV/r-based ART group (100% [15/15]) and EFV-based ART group (86.7% [13/15]).

### Tolerability and safety of artemether-lumefantrine in step 2

AL was well tolerated in the three study groups: no DAIDS grade 3 or 4 haematological or hepatic abnormalities were reported across the groups. On day 3, QTc prolongation (>450ms) was observed in 1 participant in EFV-based ART group and another in the ART-naive group but none in the LPV/r-ART group. All cases resolved by day 7.

## DISCUSSION

This study found that, when treated with a half-dose adult course of AL, individuals on EFV-based ART regimen had lower lumefantrine exposure (AUC_0-14 days_) than ART-naïve individuals while those on NVP-or LPV/r-based ART groups had higher AUC_0-14 days_. Similarly, compared to the ART-naïve group, C_max_ was lower in the EFV-based ART group, higher in the LPV/r-based ART group and similar in the NVP-based ART group. There were no differences in t_max_ across the study groups. The terminal-half life was significantly lower in the EFV-based ART group but similar in the LPV/r-or NVP-based ART groups when compared to the ART-naïve group. DAIDS grade 3 or 4 treatment-emergent thrombocytopenia and neutropenia were observed upon co-administration of AL and NVP-based ART. When treated with a standard-dose adult course of AL, there was no statistically significant difference in lumefantrine AUC_0-14 days_ between the EFV-based ART group and the ART-naïve group but those on LPV/r-based ART had higher AUC_0-14 days_ than the ART-naïve group. There were no significant differences in terminal half-life, C_max_ and t_max_ between the ART-groups and the ART-naïve group. Additionally, AL was well tolerated across all study groups.

Our finding, in both steps, of a higher lumefantrine exposure (AUC_0-14 days_) and C_max_ in the LPV/r-based ART group is consistent with what is known about ritonavir-boosted lopinavir inhibition of CYP450 enzymes (CYP3A4), resulting in higher plasma lumefantrine concentration since lumefantrine is metabolised by this enzyme entity (13, 14, 24). The therapeutic implications of this observation have been previously shown among Ugandan children who had a reduced risk of malaria treatment failure when taking lumefantrine and lopinavir-based ART compared to those not on NNRTI-based ART (15).

Unlike in step 1, where lumefantrine exposure in the EFV-based ART group was significantly lower in comparison to the ART-naïve group, overall lumefantrine exposure (AUC_0-14 days_) in step 2 was surprisingly not significantly different between the two groups. Lumefantrine concentrations in the terminal elimination phase however, were consistently lower in the EFV-based ART group compared to the ART-naïve group in both steps (Figures 1a and 1b). Since EFV is a known inducer of CYP3A4 enzymes (9), lower lumefantrine concentrations were expected in the terminal elimination phase. The difference in lumefantrine exposure in the EFV-ART/ART-naïve comparison could be as a result of the use of a parallel-group study design, which is more prone to effects of inter-individual anthropometric and genetic variations in CYP450 enzymes than in a cross-over design. Genetic polymorphisms in CYP450 enzymes are known to impact exposure of drugs metabolised by this enzyme entity (25, 26). Nevertheless, the lower lumefantrine concentrations in the elimination phase among participants on efavirenz-based ART in step 2 is consistent with previous observations (27).

There are conflicting published results on the PK interactions between AL and NVP-ART, with studies suggesting higher (16, 28), lower (18, 24) or similar (17, 19) lumefantrine exposure in those on AL and NVP-based ART compared to individuals on AL alone. This heterogeneity potentially points to genetic variations in CYP activity across HIV-malaria endemic settings. We found higher concentrations of lumefantrine in the NVP-based ART group in step 1 than in the ART naïve group, consistent with findings from an earlier study in South Africa (16) and another study conducted in Malawi and Uganda (28). There is evidence that NVP may increase exposure of other drugs metabolised by the CYP3A4 results as shown with increased C_max_ and AUC of darunavir (29) and maraviroc (30), when co-administered with nevirapine, possibly due to reduced metabolism secondary to competitive inhibition of metabolic enzymes (31) or as a result of variations in availability of proteins to transport drugs (32). Thus, the increased AUC_0-14_ days and C_max_ of lumefantrine in the NVP-based ART group could suggest reduced CYP3A4-mediated metabolism or unavailability of proteins to transport lumefantrine.

Neutropenia has been previously documented when ACTs such as artesunate-amodiaquine were administered among HIV infected children in Uganda (33). In addition, NVP is associated with granulocytopenia as a marker of hypersensitivity (34) but its role in causing thrombocytopenia has not been described. Thus, it is possible that neutropenia could occur following co-administration of NVP and lumefantrine, as a result of increased lumefantrine concentration, increased NVP concentrations or a synergistic effect of lumefantrine and NVP. In our study population, the occurrence of neutropenia across all study groups in step 1, which were not sustained at higher doses in step 2, is likely idiosyncratic. Apart from the underlying HIV infection and with the exception of those on LPV/r-based ART who took it together with zidovudine-ART, none of the participants who experienced thrombocytopenia had other baseline predisposing factors, such as low immunity (CD4 <500 cells/mm^3^) or low platelet count. Nevertheless, the data and safety monitoring board recommended against administration of a standard-dose adult course of AL with NVP due to the frequent occurrence of thrombocytopenia in addition to neutropenia in the NVP group compared to the ART-naïve group in step 1. We, therefore, were unable to investigate the effect of co-administration of a standard-dose adult course of AL and NVP on the incidence of thrombocytopenia.

Day 7 lumefantrine concentrations are considered to be one of the most important predictors of treatment outcomes following malaria treatment (35, 36). Various investigators have suggested different day 7 lumefantrine cut-offs (37–45) and in a pooled analysis, the WorldWide Antimalarial Resistance Network (WWARN) observed that day 7 lumefantrine concentrations ≥ 0.2 μg/mL (200 ng/mL) were associated with a 98% cure rate in uncomplicated malaria patients (parasitaemia <135,000/μL) (46). In step 2 of this study, although participants on EFV-ART had lower day 7 lumefantrine concentrations than ART-naïve participants and those on LPV/r-based ART had higher concentrations, the proportion achieving lumefantrine concentration ≥0.2 μg/mL was only slightly lower in the EFV-ART group but was not significantly different from the ART-naïve group. This suggests that AL is still likely to be highly efficacious in those on EFV-based ART, despite the PK interaction.

In this study, we did not assess impact of ART on plasma concentrations of the artemisinin derivatives (artemether and its metabolite, dihydroartemisinin) which have a shorter half-life and are crucial in clearing malaria parasites in the early phases of malaria treatment, because we were interested in the longer acting drug, lumefantrine, which confers protection against recrudescence following malaria infection (37, 47). Additionally, we did not quantify NVP plasma concentrations and were not able to assess any potential effect of lumefantrine on the steady-state concentration changes of NVP and subsequent impact on haematological changes. Furthermore, this study was not designed to elucidate the mechanism of interaction between lumefantrine and ART. Future studies should aim to define these mechanisms, including the role of genetic variations in CYP450 isoenzyme activity and the impact of ART on plasma concentrations of artemisinin derivatives and subsequent implication on clearance of malaria parasites among HIV-malaria co-infected individuals.

In conclusion, we confirmed that co-administration of AL with ritonavir-boosted lopinavir-based antiretroviral therapy resulted in increased lumefantrine exposure while co-administration of AL with EFV-based ART was associated with lower lumefantrine concentrations, particularly in the terminal elimination phase. Co-administration of AL and NVP-ART was associated with higher lumefantrine exposure and haematological abnormalities (thrombocytopenia and neutropenia) at half-dose adult course of AL. The therapeutic implications of these findings need to be evaluated in programmatic settings among malaria and human immunodeficiency virus co-infected individuals.

## ACKNOWLEDGEMENTS

We thank all trial participants for their participation in this study and the study team for their unwavering dedication. We would like to thank Prof Steve Ward for supporting the laboratory training of study personnel in PK assay methods. This work was supported by European & Developing Countries Clinical Trials Partnership (EDCTP) [IP.07.31060.003 to VM]. CGB acknowledges funding support from WHO-TDR as part of an EDCTP-TDR Clinical Research and Development Fellowship. The funders had no role in study design, data collection and interpretation, or the decision to submit the work for publication.

